# EvoFreq: Visualization of the Evolutionary Frequencies of Sequence and Model Data

**DOI:** 10.1101/743815

**Authors:** Chandler D. Gatenbee, Ryan O. Schenck, Rafael Bravo, Alexander R.A. Anderson

## Abstract

High throughput sequence data has provided in depth means of molecular characterization of populations. When recorded at numerous time steps, such data can reveal the evolutionary dynamics of the population under study by tracking the changes in genotype frequencies over time. This necessitates a simple and flexible means of visualizing an increasingly complex set of data. Here we offer EvoFreq as a comprehensive tool set to visualize the evolutionary and population frequency dynamics of clones at a single point in time or as population frequencies over time using a variety of informative methods. EvoFreq expands substantially on previous means of visualizing the clonal, temporal dynamics and offers users a range of options for displaying their sequence or model data. EvoFreq, implemented in R with robust user options and few dependencies, offers a high-throughput means of quickly building, and interrogating the temporal dynamics of hereditary information across many systems. EvoFreq is freely available via https://github.com/MathOnco/EvoFreq.

## Background

Changes in genotype frequencies are often visualized using Muller plots, wherein each polygon represents a genotype (clone), and the thickness of the polygon indicates either the number of individuals with that genotype, or the frequency of the genotype in the total population at each time point. Nesting of genotypes represents evolutionary relationships, i.e. one genotype emerging from within another genotype’s polygon indicates that the former was created by an individual with the latter genotype. Muller plots thus provide an excellent way to visualize how the genetic composition of a population changes over time. As these changes are governed by mutation, selection, drift, and gene flow (evolutionary forces), the visualization of these dynamics facilitates an understanding of which forces are dominating the population.

One example where this form of data visualization has proven insightful is in understanding tumor evolution and subclonal compositions from both mechanistic models and bulk or multi-region sequencing. The hereditary nature of mutations in somatic tissue and cancers allows us to recapitulate, the temporal dynamics associated with mutational events through subclonal reconstruction. To date, two visualization packages exist in R to visualize this temporal data fishplot [5] and ggmuller [7]. We constructed an alternative library that includes the features of these two libraries while expanding functionalities (see Table 1). EvoFreq offers users ease of implementation and a way to generate many visualizations for assessing the temporal dynamics of clonal <u>Evo</u>lutionary Frequencies. EvoFreq is available at https://github.com/MathOnco/EvoFreq.

**Table 1.** Comparison of the features within the currently available packages for plotting evolutionary dynamics. EvoFreq is capable of visualizing the evolutionary dynamics from ClonEvol, PhyloWGS, and CALDER.

|  | ggmuller | fishplot | EvoFreq |
| --- | --- | --- | --- |
| Frequency data |  | ✓ | ✓ |
| Size data | ✓ |  | ✓ |
| Long format | ✓ |  | ✓ |
| Wide format |  | ✓ | ✓ |
| Smoothing |  | ✓ | ✓ |
| Uses ggplot | ✓ |  | ✓ |
| Animations |  |  | ✓ |
| Create graphs |  |  | ✓ |
| Integration with subclonal reconstruction software |  | ✓ | ✓ |

## Implementation and Results

EvoFreq is built on top of the leading visualization library in R, ggplot2 [13]. Input data can be either clone sizes or frequencies, and in long or wide formats. Given such data, EvoFreq can create Muller plots and/or dendrograms, revealing the clonal dynamics over time. Additional customizations include the ability to color clones based on a user defined attribute (such as fitness), provide custom colors, and label polygons. Due to EvoFreq’s utilization of ggplot2 as the primary, underlying library, further customization to EvoFreq’s plots is possible. Using the optional dependency [8], users may also create animations of evolving Muller plots and “growing” dendrograms.

EvoFreq is capable of visualizing frequency dynamics at a single point in time, as a phylogenetic tree representation, a graph representation, or as a frequency plot over time similar to FishPlot and ggMuller, but with extended capabilities.

At the core of EvoFreq is the ability to visualize relational data structures over time, whether this is from simulations or inferred from data. Here we illustrate the usefullness of generating these results using EvoFreq. Our first example utilizes data generated by West *et al.* which quantifies how spatial constraints alter the evolutionary trajectory of a single tumor using a passenger-driver model [10]. EvoFreq was used extensively within this publication and a subset of this data is shown in Figure 1. This figure highlights how the user can manually or automatically add labels to provide informative details of clones, a particularly useful feature of EvoFreq.

**Figure 1.**
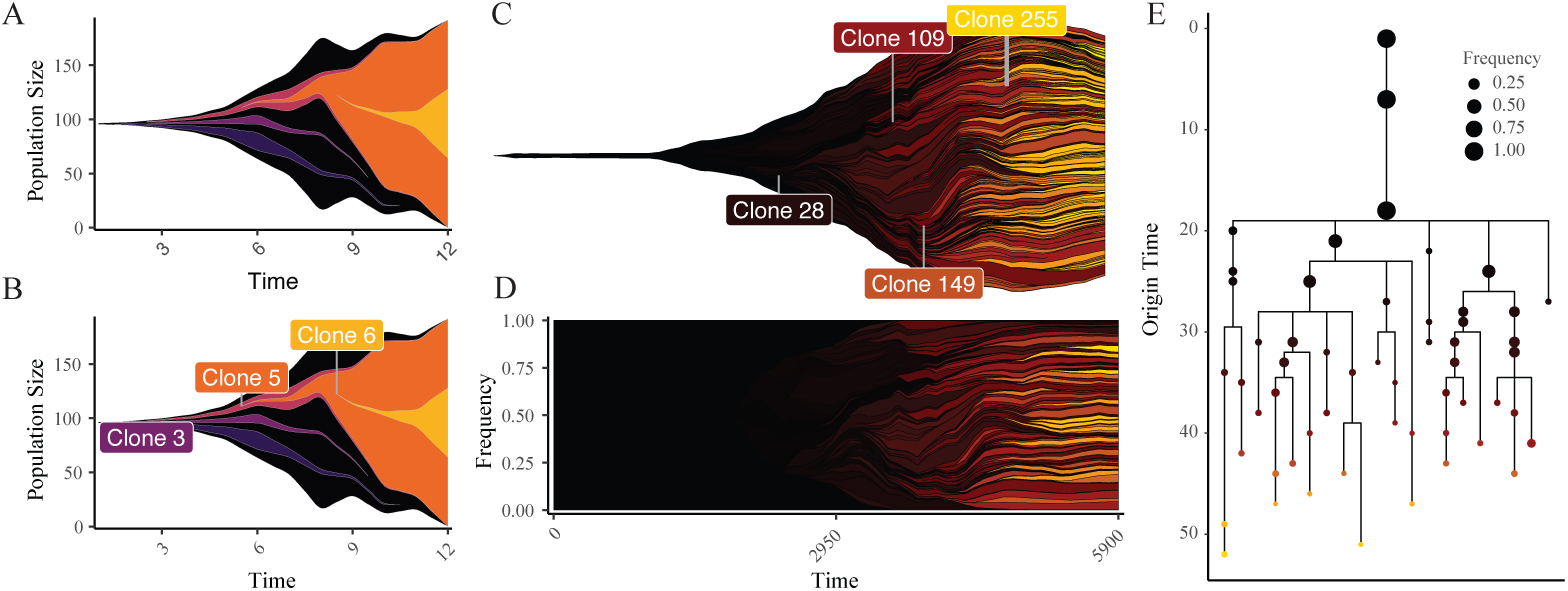
EvoFreq is a comprehensive and flexible R package for the visualization of longitudinal data. **A** and **B** show an EvoFreq plot for one of the provided datasets with **B** and without **A** the function to add labels. For more complicated data EvoFreq provides a powerful means to quickly filter data (**C** and **D** thresholded at 0.2 frequency), color by an attribute (driver strength), and visualize dynamics as a frequencies rather than population size. Using a dendogram styled to ensure that origin time is easily conveyed (termed here as an EvoGram), a more quantitative view is provided **E**.

Second, we analyze sequence data using three different clonal reconstruction tools from 15 initial engraftments and serial propagation of primary and metastatic breast cancers [3] and visualize these results using EvoFreq. We applied ClonEvol [1], PhyloWGS [2], and CALDER [6] to infer the clonal dynamics from each of the longitudinal xenografts (select inferences are illustrated in Figure 2). Initial processing and reformatting of the somatic single nucleotide variant (SNV), copy-number alteration (CNA), and loss of heterozygosity (LOH) data from whole-genome shotgun sequences (WGSS) and Affymetrix SNP Array 6.0 was carried out to extract read data using custom python scripts and prepare inputs for PhyloWGS and CALDER. Each of these tools requires different pre-processing and infers subclonal reconstructions in different ways. We have incorporated functions within EvoFreq to parse outputs of each of these tools to visualize inferred clonal dynamics using EvoFreq. When the output provides numerous solutions for subclonal reconstructions, the user has an option of returning one or all of these solutions. Examples of this process have also been provided within EvoFreq’s documentation.

**Figure 2.**
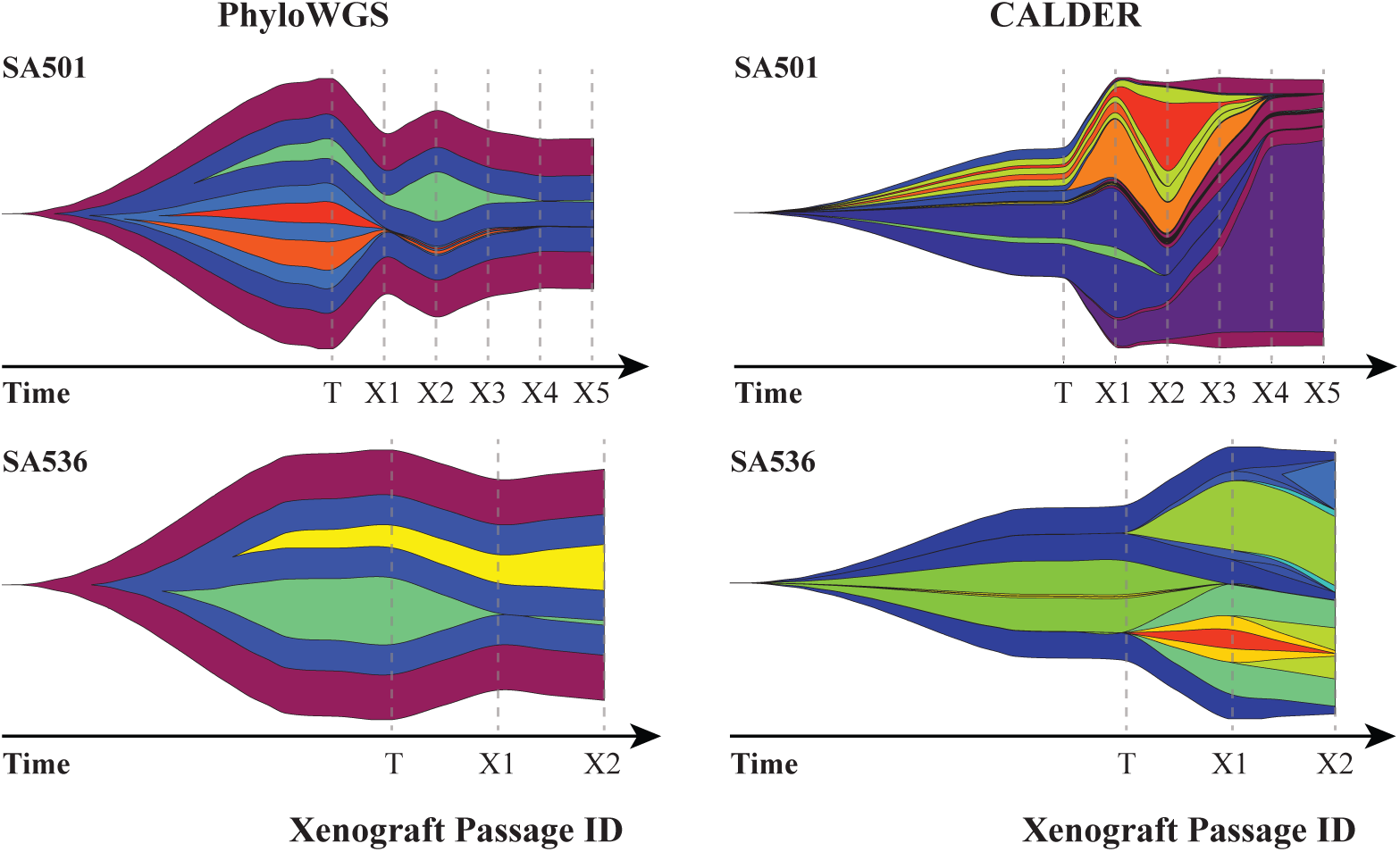
EvoFreq can be easily used to visualize outputs from CloneEvol, PhyloWGS, and CALDER. Data parsing functions are included within EvoFreq to rapidly visualize subclonal reconstructions. Each column above illustrates a method of subclonal reconstruction for two separate human breast cancer xenograftments from Eirew *et al.* [3] for PhyloWGS (left) and CALDER (right). Originating tumors (T) and their subsequent xenograft passages (X1, X2, etc.) are shown for SA501 (top) and SA536 (bottom).

## Conclusions

We present EvoFreq as a versatile library capable of, generating publication and presentation ready images as well as video for, visualizing clonal frequencies over time. EvoFreq’s design allows for broad access to all users through robust support for input data and input validation functions. EvoFreq can be used for all relational data structures providing many different visualizations from one user-facing library, making it applicable in a number of fields. EvoFreq has currently been adopted by a number of research groups, focused on cancer genomics and mechanistic modelling, and has been used in more than four studies already [4, 9, 11, 12]. All source code, read me, and issue support is available at https://github.com/MathOnco/EvoFreq.

## Availability and requirements

**Project name**: EvoFreq

**Project home page**: https://github.com/MathOnco/EvoFreq

**Operating system**: Operating system independent.

**Programming languages**: R

**Other requirements**: ggplot2.

**License**: GNU GPLv3

**Any restrictions to use by non-academics**: None

## Author’s contributions

CDG and ROS wrote the code base and packaged EvoFreq for distribution and prepared the manuscript. RRB assisted in the development of algorithms needed for graphical representation of frequencies. ARAA provided guidance on code development, oversaw all work efforts and provided funding. All authors read, edited, and approved the manuscript.

## Availability of data and materials

Not applicable.

## Ethics approval and consent to participate

Not applicable.

## Consent for publication

Not applicable.

